# Stable expression of *Helicobacter pylori cagA* oncoprotein in brinjal

**DOI:** 10.1101/2023.05.12.540570

**Authors:** Mohammad Javad Mehran, Rambod Barzigar, Basaralu Yadurappa Sathish Kumar, Nanjundappa Haraprasad, Bashasab Fakrudin, Sayan Paul, Rajesh Kumar Ramasamy, Sudhakar Malla

## Abstract

*Helicobacter pylori* is closely connected to upper gastrointestinal tract diseases including gastric cancer. Transgenic plants are found to be successful in expressing the bacterial antigens, which could elicit an immune response when consumed. The Cytotoxicity-associated immunodominant antigen protein (*cagA*) of *H. pylori* is kindred with pathogenicity and cancer risk. We expressed the *cagA* transgenically in the brinjal. We amplified the *cagA* gene from *H. pylori* strain 26695 chromosomal DNA and transformed it into brinjal callus derived from leaf explants using the pBI121 expression vector. The stable expression and accumulation of the recombinant *cagA* gene were confirmed by using quantitative real-time PCR, western blot analysis and ELISA. The RT-PCR, western blot and ELISA showed stable expression of *cagA* gene in the transgenic lines B3, B5, B11, B17 and B21. Among them, B11 and B17 samples showed higher expression of the *cagA* compared to the other samples. Besides, the immunohistochemistry assay showed the abundant expression of *cagA* protein in the parenchymal regions of the transgenic plants. Out of the 52 plants, a set five plants were found to be positive for *cagA* expression. Our experimental outcomes can be used further to design the vaccines against *H. pylori* from the transgenic brinjal plants.

## Introduction

*Helicobacter pylori*, a stomach colonizing bacterium is widely and infectious (Ohnishi et al., 2008). It is a chronic bacterial infection that is the primary reason for spawning gastric cancer in humans (Nešić et al., 2014). It is a human gastric pathogen, infecting more than half of the world’s population (Hooi et al., 2017), which leads to chronic infection in the absence of antibiotic treatment (Malfertheiner et al., 2017). Gastric cancer is thought to be frequently caused by chronic *H. pylori* infection (Fallone et al., 2016). Around the world, more than 89% of gastric cancer cases are found to be related to *H. pylori* infection (Josephson and Skole, 2018; Turabi et al., 2022). In 1994, the WHO International Agency for Research on Cancer (IARC) confirmed *H. pylori* as a Group I carcinogen and its eradication is very important to prevent gastric cancer among both developed and developing nations (Hooi et al., 2017; Chatterjee et al., 2021). Increasing antibiotic resistance is also responsible for the failure in prophylaxis against this pathogen, which is again leading to a downfall in the prognosis of *H. pylori* eradication (BARZIGAR et al., 2020). In Europe alone, the traditional usage of triple therapy was stopped and replaced with quadruple therapy (Graham and Shiotani, 2008). However, due to the lack of proper guidelines in quadruple therapy, antibiotics are suggested randomly just to reduce the rate of antibiotic resistance.

People infected with *H. pylori cagA*-positive strains are said to be more prone to gastric cancer. *cagA* (cytotoxin-associated gene A) is a 120–145kDa protein that is responsible for the virulence of the strain to cause human cancer (Barzigar et al., 2021). This *cagA* protein encoded by the *cagA* gene is dispatched into epithelial cells of the stomach through bacterial type IV secretion which then encounters tyrosine phosphorylation. This in turn flirts as a non-physiological protein interacting with many signaling molecules like pro-oncogenic phosphatase and polarity-regulating kinase (PAR1) (Backert et al., 2011). This manipulation conciliated by intracellular signaling not only boosts neoplastic malformations but also destroys the epithelial lining of stomach cells (Hatakeyama, 2017).

The most effective way to control the infection of this bacterium is vaccination (Pan et al., 2018). As urease subunit B (*ureB*) gene was found to be more protective than urea it was considered as a potential antigen against *H. pylori*. Cost is one of the major barriers to produce new vaccines for populations that need them urgently. It has been estimated that downstream processing and purification of recombinant proteins represent 80% to 90% of the cost of recombinant pharmaceuticals (Sabalza et al., 2014). Several factors such as plant pathogens, phenolic compounds, and secondary metabolites could affect the difficulty of purification at an industrial level (Abdoli-Nasab et al., 2013). Plant-derived recombinant proteins will be inexpensive in production with good product quality and free from other animal viruses. The development of new vaccines and drugs is one of the vital goals in molecular farming especially in developing countries (Soria-Guerra et al., 2007). One of the advantages of plant expression systems is the production of edible vaccines using leafy plants, fruits or vegetables. The oral-based plant delivery system would eliminate the purification process, sterile injection hazards, health care workers, and also cold storage (Streatfield and Howard, 2003). Oral delivery promotes the induction of systemic and mucosal immunity; additionally, bioencapsulation by cellulosic plant cell walls protects proteins from digestion (Tiwari et al., 2009).

Brinjal or aubergine, commonly called eggplant, is notably known as the common man’s vegetable in India and many Asia and Latin American countries (Wei et al., 2019). It is also crowned as a poor man’s vegetable owing to its accepted usage among people and farmers with low income, and it is often crowned as the ‘King of Vegetables’ (Meyer et al., 2012). It is very often used in ayurvedic formulations to cure diabetes, blood pressure and obesity. The brinjal plant belongs to the family *Solanaceae*, and it is cultivated in many parts of the world. The brinjal has the ability to respond well in tissue culture, particularly plant regeneration from cultured seedling explants (leaf, cotyledon and hypocotyl). Brinjal also responds well to *Agrobacterium*-mediated transformation with both cointegrate and binary vectors (Iannacone et al., 1997). A number of plant systems including *Arabidopsis*, rice, carrot and tomato were used as recombinant models for the expression of various genes including potential pharmaceuticals (Saini and Kaushik, 2019). However, brinjal is not explored to a great extent as a potential genetic engineering model in this field and only limited reports describing the use of eggplant in molecular farming (Mehran et al.; Mehran et al., 2021). So, we have decided to use brinjal for our study. To probe for the possible usage of plants to produce *H. pylori cagA*, efforts have been made in this study to express the clone of *cagA* gene from *H. pylori* strain 26695 in the brinjal, *Solanum melongena L. Cultivar* Arka Keshav. Using brinjal plants, we demonstrated the expression of the *cagA* gene. Our results insinuate that further studies could throw more light in bringing out possible vaccines against *H. pylori* from the brinjal plant.

## Materials and Methods

### Amplification of cagA gene from H. pylori 26695

*cagA* forward and reverse primer pairs were used for PCR amplification. To clone the DNA corresponding to ∼1700bp, synthetic oligonucleotide primers determining the sequence of *H. pylori* were constructed.

In a total volume of 25μl, the amplification reaction contains roughly 50ng of total gDNA as template, Hotstart Taq DNA polymerase (1U), dNTPs (0.2mM each), 1 PCR buffer, MgCl2 (3mM), and primers (10pmol of each primer). Initial denaturation at 94°C for 10 minutes was followed by 35 cycles (94°C, 30sec; 55°C, 30sec; 72°C 2min) and a final extension at 72°C for 10 minutes. After that, the produced products were separated on a 1% agarose gel and the results were determined using a UV transilluminator (Paul et al., 2018; Ramasamy Rajesh Kumar, 2020; Saravanakumari et al., 2020; Paul et al., 2022b). The DNA fragments were purified from the agarose gel using the Genei clean purification technique (Genei Gel Extraction Kit). According to the instructions, the Miniprep Kit (Genei) has been used to obtain pure super-coiled plasmid DNA with excellent yield (35μl) (Arumugaperumal et al., 2020; Garcia Jr et al., 2020; Kumar et al., 2020).

### Cloning of cagA gene in pBI121

We cloned the *cagA* gene, which is around 1700bp long, in order to utilise the plants to stimulate an immunological response in *H. pylori* infected individuals. After that, the gene was amplified, and the pure product (1%) from an agarose gel was cleaved with *BamHI* and *SacI* (37^0^C for 2hr). The restriction enzymes were also used to digest the vector. Using the Genei gel extraction kit, the bands were removed and cleaned. The linearized vector and *cagA* product then were ligated overnight at 16^0^C using T4 DNA ligase, then transformed into the DH5α strain and verified by sequencing (Eurofins, Bangalore).

### Preparation of competent cells and transformation

All *E. coli* cells are made competent using the calcium chloride approach (Chang et al. 2017), then streaked on LB-agar plates without antibiotics and incubated for 12 hours at 37°C. A single colony was injected into 10mL of LB medium and cultured at 37°C until the culture attained an optical density of 0.4-0.5 at 660nm (3-4hr). The cells were cooled on ice for 30 minutes before being extracted in autoclaved tubes using an SS 34 rotor (Sorvall refrigerated centrifuge) for 15minutes at 4°C (Evolution RC). The cell pellets were resuspended in 30ml of filter-sterilized ice-cold acid salt buffer (100mM CaCl_2_, 70mM MnCl_2_, and 40mM sodium acetate, pH 5.2-5.5) and incubated for 45 minutes on ice.

The pellet was dissolved in 1/25^th^ volume of bacterial suspension in acid salt buffer containing 20% glycerol after being spun at 3000rpm for 15 minutes at 4°C (cell pellet from 90 ml culture was suspended in 4.5ml of buffer). 50µl aliquots were prepared and kept at −70°C until needed. By incubating competent DH5α *E. coli* cells on ice for 30 minutes and then heat shock at 37°C for 5 minutes, the pBI121 vector harboring the *cagA* gene was transformed. The *E. coli* cells were then cultured on LB agar plates with antibiotics (Kanamycin 20µg/ml). The plates were then incubated for 12 hours at 37 degrees Celsius, and bacteria were tested for pBI121 insertion.

### Isolation of plasmid DNA

Single isolated colony was inoculated in 5ml of antibiotic-laced LB media and cultured for 12 hours on a shaker incubator at 37°C (120rpm). The alkaline lysis procedure was used to isolate the plasmid DNA (Sambrook and Russell, 2006). The pellet from a 5ml overnight culture was spun at 6000g for 5 minutes at 4°C and resuspended in 200μl of prechilled solution I. 200μl of freshly made solution II were added and carefully mixed by tipping the container upside down. After solution II, 200μl of solution III was poured and carefully mixed, then maintained on ice for 10 minutes. The cell lysate was spun at 8000g for 10 minutes at 4°C after incubation. An equal volume of phenol, chloroform, and isoamylalcohol mixture (25:24:1; v/v) was added to the supernatant and well mixed.

The contents then were spun at 8000g for 10 minutes at 4 °C before being mixed with an equivalent amount of pre-chilled isopropanol. The DNA then was pelleted at 8000g for 10 minutes at 4°C before being reconstituted in 10mM TE buffer. On a 0.8 percent agarose gel electrophoresis, the purity of the recovered plasmid DNA was verified. Restriction digestion and DNA sequencing were used to confirm positive clones.

### Callus induction

Eggplant seeds of the variety Arka Keshav were surface sterilised using sodium hypochlorite (6%) plus water in a 1:1 ratio, then washed 3-4 times with sterile distilled water. About 20 seeds were inoculated on Petri plates with N6 callus induction medium (pH 5.75) containing 10,000mg/l Myoinositol, 200mg/l Glycine, 50mg/l Thiamine HCl, 50mg/l Pyridoxine HCl, and 50mg/l Nicotinic acid. After autoclaving the previously mentioned contents, 10M BAP and 1M NAA were added to the medium. The medium was solidified with 0.8 percent agar. After that, the Petri plates were incubated in the dark at 252°C. The calli were then extracted from the growing seeds and subcultured on a new callus induction medium. Explants in our investigation included median sections of cotyledonary leaves plus crosswise cut hypocotyl segments.

### Agrobacterium-mediated transformation

With minor modifications, the calli produced from the seeds were employed in *Agrobacterium* (EHA105)-mediated transformation as described by Patel et al., 2013 (Patel et al., 2013). Electroporation was used to introduce the vectors into *Agrobacterium*. In a nutshell, 5 litres of plasmid DNA were put to 45 litres of cells and gently mixed. The contents are electroporated after being put to a prechilled cuvette. The vector constructs were implanted into 5ml of YEP (yeast-extract-mannitol medium; pH 7.0) medium (containing 20mg/l Rifampicin and 50mg/l Kanamycin) and cultured at 28°C in a shaking incubator (200rpm) overnight. Overnight culture broth was added to 45ml of infection medium (MS baseline media with Thiamine HCl [1mg/l], Myoinositol [250mg/l], Casein hydrolysate [1.0g/l], Proline [690mg/l], Glucose [30g/l], 2,4-D [5.0mg/l], and Acetosyringone [200M]) at pH 5.2 and incubated at 28°C. *Agrobacterium* colony PCR was used to validate the insert *cagA*.

### Isolation of genomic DNA from leaves

Genomic DNA was isolated from the leaves by the CTAB method as described by Li, Z *et al*., 2020 (Li et al., 2020). In brief, about 100mg of leaf tissue was homogenized in a micro pestle with 0.5ml of extraction buffer (2% CTAB, 1.4M NaCl, 20mM EDTA, 10mM Tris HCl; pH 8.0) with a 10µ1 of beta-mercaptoethanol. The contents were mixed thoroughly and incubated for 30-40min at 65°C in a water bath, and added with equal volumes of chloroform: isoamyl alcohol (24:1) following incubation. The contents were then centrifuged at 5000rpm for 15min at room temperature and added with 1/10^th^ volume of 5M NaCl and added with 0.8volumes of ice-cold isopropanol to precipitate the DNA. The DNA obtained was then pelleted at 8000rpm for 10min and washed with 70% ethanol. Following washing, the pellet was resuspended in 100µl of TE buffer. 5µl of RNase (l0mg/ml) was added to the DNA and stored at −20° C following incubation at 37°C for 1 hour.

### PCR amplification of cagA

Real-time PCR as well as western blot were used on total RNA separated from leaves of wild-type and transgenic plants to measure the expression level of *cagA* mRNA. In the control group, no bands were seen. As described in the preceding chapter, PCR amplification was carried out. The template was made using genomic DNA extracted from the plants. To validate the *cagA* gene insert, the products then were tested on a 1% agarose gel. The Primer 3 programme (version 4.13) was used to create all primers utilised in this work (Table 1), which were predicated on mRNA sequences deposited in GenBank (Untergasser et al., 2012). Primers were crosschecked for specificity using BLASTN algorithm (Chinnadurai et al., 2018; Alaguponniah et al., 2020; Kumavath et al., 2021; Miao et al., 2022). They may amplify genomic DNA and also cDNA since each primer set is placed on the same exon of the gene. As a result, DNase treatment was used to guarantee that the RNA samples were devoid of DNA. The amplification of cDNA using the chosen primers was verified using conventional RT-PCR tests followed by gel electrophoresis (Bleve et al., 2003; Rana et al., 2022b; Rana et al., 2022a).

### RNA extraction and cDNA synthesis

The plant leaves are flash-frozen with liquid nitrogen and preserved at –80°C after steady transformation of the Brinjal plant expression. The whole leaves were pounded into a fine powder in liquid nitrogen using a mortar and pestle. The RNeasy Plant Mini Kit (Genei Laboratories Pvt Ltd) was used to extract total RNA from 100mg of ground tissue according to the manufacturer’s instructions. With the M-MuLV RT-PCR Kit, the cDNA was generated as reported by Ramalho *et al* 2004 (Ramalho et al., 2004). As a starting material, around 2µg of the RNA produced in the preceding phase was employed. We utilised around 1.02 litres of total RNA, random primers, and 1µl of RT enzyme because the RNA concentration was 40µg/ml.

Following the initial incubation, the contents were incubated at 25°C for 10 minutes and then at 70°C for 45 minutes. The generated cDNA was utilised to amplify the entire *cagA* gene as well as for qPCR expression experiments. The *cagA* was successfully amplified to determine the amount of *cagA* mRNA expression in the tissues. The amplification was carried out as per the protocols mentioned by Green *et al.,* 2019 (Green and Sambrook, 2019). To validate the *cagA* gene insert, the products were tested on a 1.2 percent agarose gel.

### Real-time PCR

On cDNA templates, a quantitative RT-PCR experiment was performed using the SYBR Green PCR master mix (Bio-Rad) as per the manufacturer’s protocols (BioRad) using the designed oligonucleotide primers given in table 1 (Schmidt and Delaney, 2010). A 25µl reaction including 2.5µl of 10x PCR buffer with SyBr green, 2.5µl of 10mM dNTPs, 10pmoles/l of each primer, 10.4µl of PCR water, 0.1µl of 5U Taq polymerase (Genei), and 2µl of cDNA was used to amplify the *cagA* and housekeeping genes (Actin) (Expósito-Rodríguez et al., 2008; Paul et al., 2021b; Paul et al., 2023; Ponesakki et al., 2023).

Initial denaturation of cDNA was performed at 94°C for 10 minutes, followed by 35 cycles of amplification at 94°C for 30 seconds, 55°C for 30 seconds, and 72°C for 30 seconds. At the end of the run, a last extension at 72°C for 10 minutes was performed. Duplicates were done on all of the samples, and samples without cDNA served as a negative control. The output of RT-PCR was analyzed using the CFX Maestro Software and the fold change values were calculated using the ΔΔCT and 2-ΔΔCT formula (Paul et al., 2021a; Tatta et al., 2023). The unpaired t-test was used to determine the statistical significance and the P value ≤ 0.05 was considered as statistically significant for our study (Livak and Schmittgen, 2001; Paul et al., 2022a).

### Protein extraction and western blot

Using a mortar and pestle, the leaf samples (both transgenic and control) were thoroughly pulverised in liquid N2. 0.2gm tissue powder is reconstituted in 2ml cold acetone and vortexed for 30 seconds (Wang et al., 2003). Centrifuging at 10000g for 3 minutes at 40C, the pellet was washed with acetone. The pellet was then moved to a mortar and finely powdered with quartz sand. The powder was washed 3–4 times with ice-cold 10% TCA before being mixed with cold 80 percent acetone. The pellet was maintained at room temperature after centrifugation for extraction. The study employed a slightly modified version of the phenol extraction technique (Wang et al., 2003).

In a nutshell, around 0.1gm of powder was reconstituted in 0.8ml phenol (Tris-buffered, pH 8.0) and 0.8ml SDS buffer (sucrose 30 percent, SDS 2 percent, 0.1M Tris-HCl, pH 8.0, 5 percent 2-mercaptoethanol). The components are well mixed for 30 seconds before being centrifuged at 10000g for 3 minutes. To precipitate the proteins, the top phenol phase was recovered in new tubes and mixed with 5 volumes of cold methanol and 0.1M ammonium acetate. After that, the proteins were centrifuged at 10,000g for 5 minutes before being rinsed with methanolic ammonium acetate and 80% acetone. After drying the pellet, it was reconstituted in 2-DE rehydration solution (8M urea, 4% CHAPS, 2% IPG buffer, 20mM dithiothreitol). Using bovine serum albumin as a reference, the proteins were measured using a protein assay (Genei, Bangalore).

With minor adjustments, the expression levels of transgenic plants encoding *cagA* were measured by western blotting. SDS–polyacrylamide gel electrophoresis (4.75 percent stacking and 12 percent resolving gel with 12 percent glycerol) was used to separate soluble proteins, which were then transferred to Nylon membranes (Genei, Bangalore). The resolving buffer (Tris-HCl, pH 8.8) was increased to a final concentration of 0.75M. The protein samples were denatured at 95°C for 3 minutes before being resolved at constant 200V. After blocking with 3 percent bovine serum albumin in TBST buffer (20mM Tris-HCl, pH 7.6, 0.8 percent NaCl, 0.1 percent Tween20), the blots were hybridized for about 1 hour at room temperature with affinity-purified Biotin conjugated anti *cagA Helicobacter pylori* antibody (ThermoFischer, Biotin dilution 1:500). The investigation employed the secondary antibody streptavidin (1:1000) which was identified using a western blotting detection kit (Genei, Bangalore). The study employed beta-actin with a molecular weight of ∼42KDa.

### Immunohistochemistry assay

Leaf samples were taken from both transgenic and control plants and implanted using the Technovit 7100 Embedding Kit (Heraeaus Kulzer, Wehrheim, Germany). The immunohistochemistry test was carried out according to Bo Hu et al., 2012’s procedure (Hu et al. 2012). 5mm thick transverse slices were cut and dried on slides using a microtome. At room temperature, the slices were treated in H2O2 (3%) for roughly 15 minutes to prevent any peroxidase activity. The sections were rinsed three times with distilled water and masked with BSA after incubation (10 percent). After that, the slides were treated at 4°C overnight with affinity-purified Biotin conjugated anti *cagA Helicobacter pylori* antibody (ThermoFischer, Biotin dilution 1:2000). The slides were then washed 3-4 times in PBS before being treated for 30 minutes at room temperature with a biotin-labeled goat anti-rat IgG antibody. After washing with PBS, the slides were incubated for 30 minutes at room temperature with SABC (streptavidin and biotinylated horseradish peroxidase complex (SABC) reagent (Sigma-Aldrich, USA). The slides were then stained with the AEC agent kit (HiMedia Ltd) at 370C after thorough washing with PBS (0.02% Tween 20 and PBS). After that, the slides were cleaned in distilled water and inspected with an Olympus IX-70 microscope. Negative controls were slides treated with PBS/BSA but not with primary antibodies.

### ELISA

The *cagA* transgenic brinjal leaves extracts were diluted as 1:5 and 1:10 in sodium bicarbonate (pH 9.8) and coated to the wells of the ELISA plate by adding 4.5–5 mg/mL concentration of antigen solution and incubating at 37 °C for 2 h, followed by three washes with washing buffer (0.1 percent Tween 20 in PBS) for 5 min each time (Barzigar et al., 2022). The plate was blocked for 90 minutes with 2 percent skim milk, then cleaned three times with washing solution. The plate was then treated for 1 hour at room temperature with anti *cagA Helicobacter pylori* antibody (ThermoFischer, Biotin dilution 1:500). Wells were rinsed again with the washing solution after incubation, and of streptavidin-HRP conjugate (1:10,000) was added to each well and incubated for 30min. The reaction was terminated with 0.5 M HCL, and the absorbance was measured at 450 nm with a Bio-Rad microplate reader (Nayeri and Anbuhi, 2019). As a negative control, total protein was isolated from non-transgenic brinjal leaves. All of the trials were carried out in threes.

## Results

### Amplification and cloning of cagA gene into pBI121

The PCR amplification with *cagA* primers resulted in a positive amplicon of around 1700bp as confirmed by the bands [Figure 1]. The isolated plasmids run on an agarose gel revealed two products after digestion with restriction enzymes, one of which contained the *cagA* gene product at around 1700bp. The acquired nucleotide sequences were then compared to their GenBank counterparts. Using BLAST, the sequence was discovered to have a “perfect” match (97.93 percent similarity) with sequences of respective corresponding gene *cagA* from GenBank (GenBank® accession no. CP026326.1). This variation could be widespread among strains.

**Figure 1:**
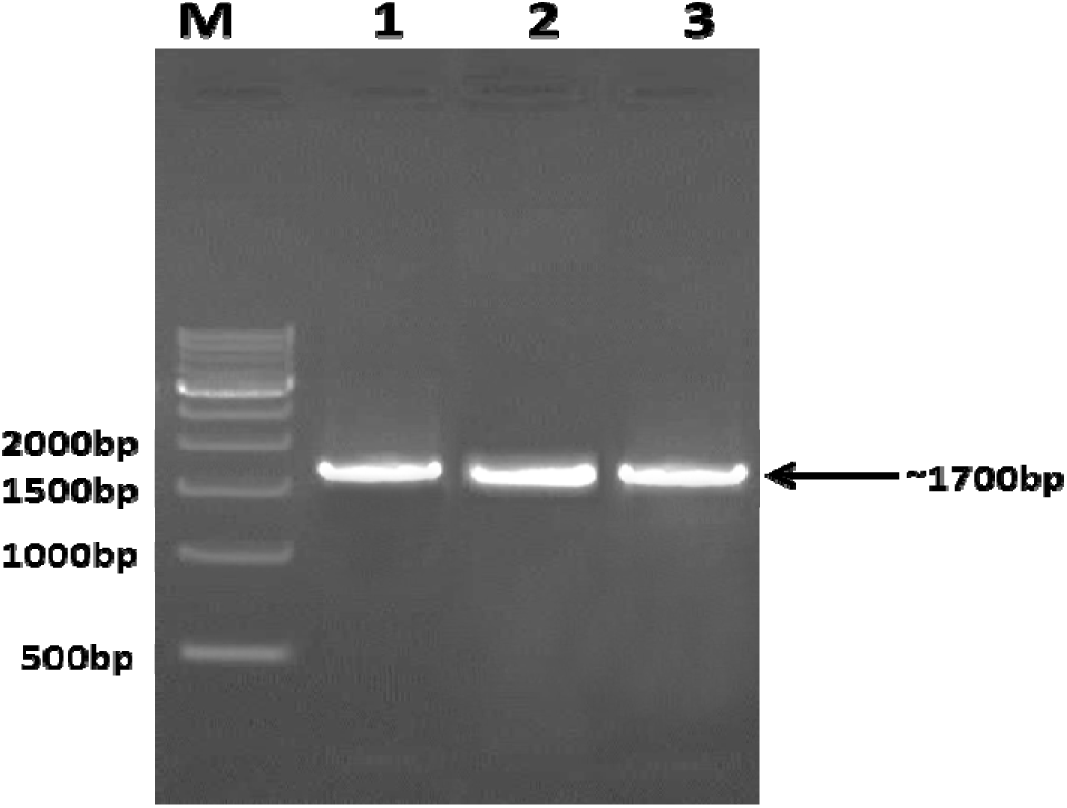
PCR amplification of *cagA* gene *H.pylori* 26695 by specific primers. Image showing the *cagA* gene PCR amplification on 1% agarose gel. Lane description: *cagA* gene product lane 1, 2, 3 and molecular-weight ladder (M). Image as viewed on Gel Dock.

### Agrobacterium-mediated genetic transformation

*Agrobacterium*-mediated transformation was used to rejuvenate 52 separate brinjal plants [Figure 2]. The positive expression of *cagA* was also verified by *Agrobacterium* colony PCR. In the gel, there were bands with a size of 1700bp. PCR amplification and immunoblot analysis were used to check for *cagA* expression. The genomic DNA of all five transgenics, as well as the cDNA of the transgenic plants, yielded a PCR result of the predicted size (1700bp), indicating that the gene is expressed positively [Figure 3]. The control (independent non-transgenic) genomic DNA and cDNA revealed no band with negative amplification, as predicted. There is no evidence of non-specificity, indicating that the DNA is pure and specific.

**Figure 2:**
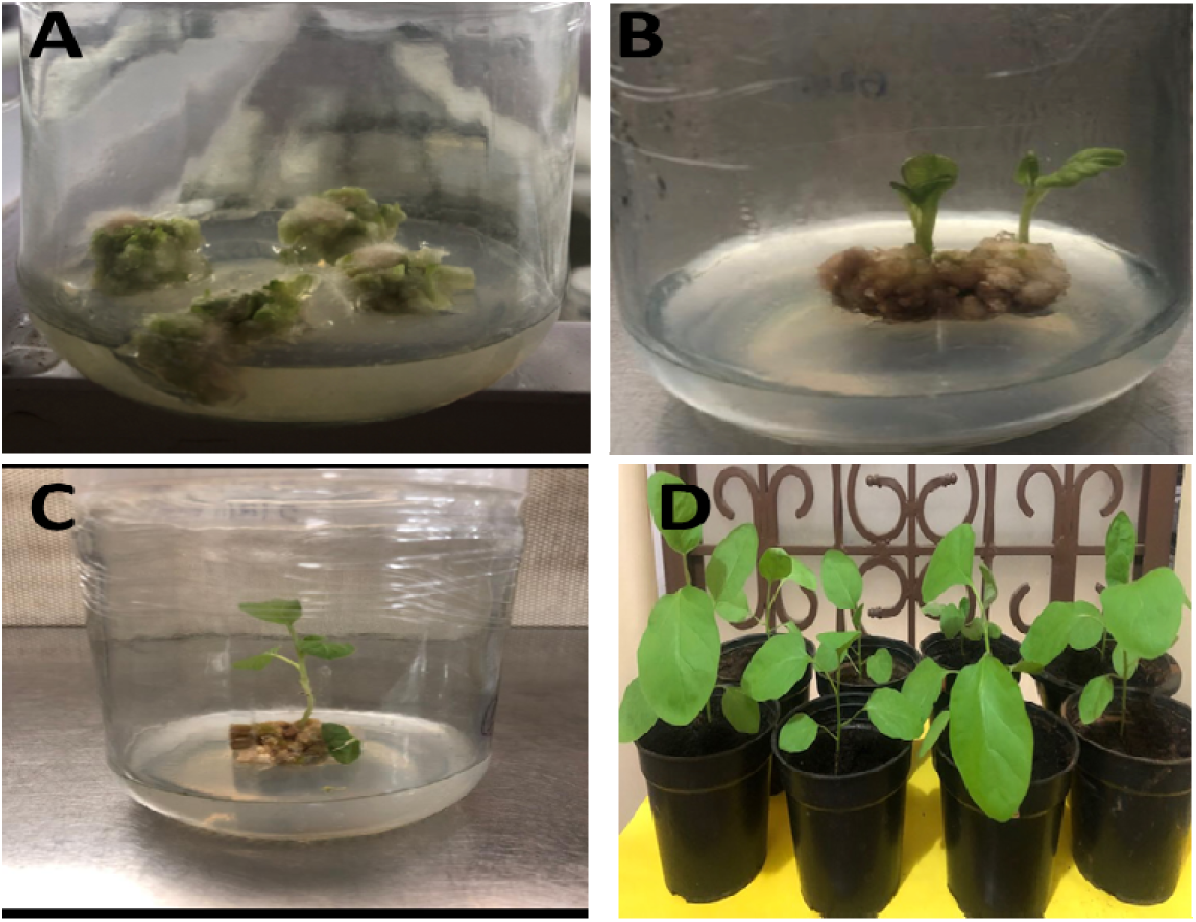
Production of transgenic brinjal plants by *Agrobacterium-*mediated Transformation. Images showing the development of transgenic brinjal *cv* Arka Keshav, A: Callus initiation from *Agrobacterium* transformed leaf discs on selection medium, B, C: Kanamycin-resistant transgenic shoots cultured on MS medium for rooting, D: Potted transgenic brinjals grown in the soil.

**Figure 3:**
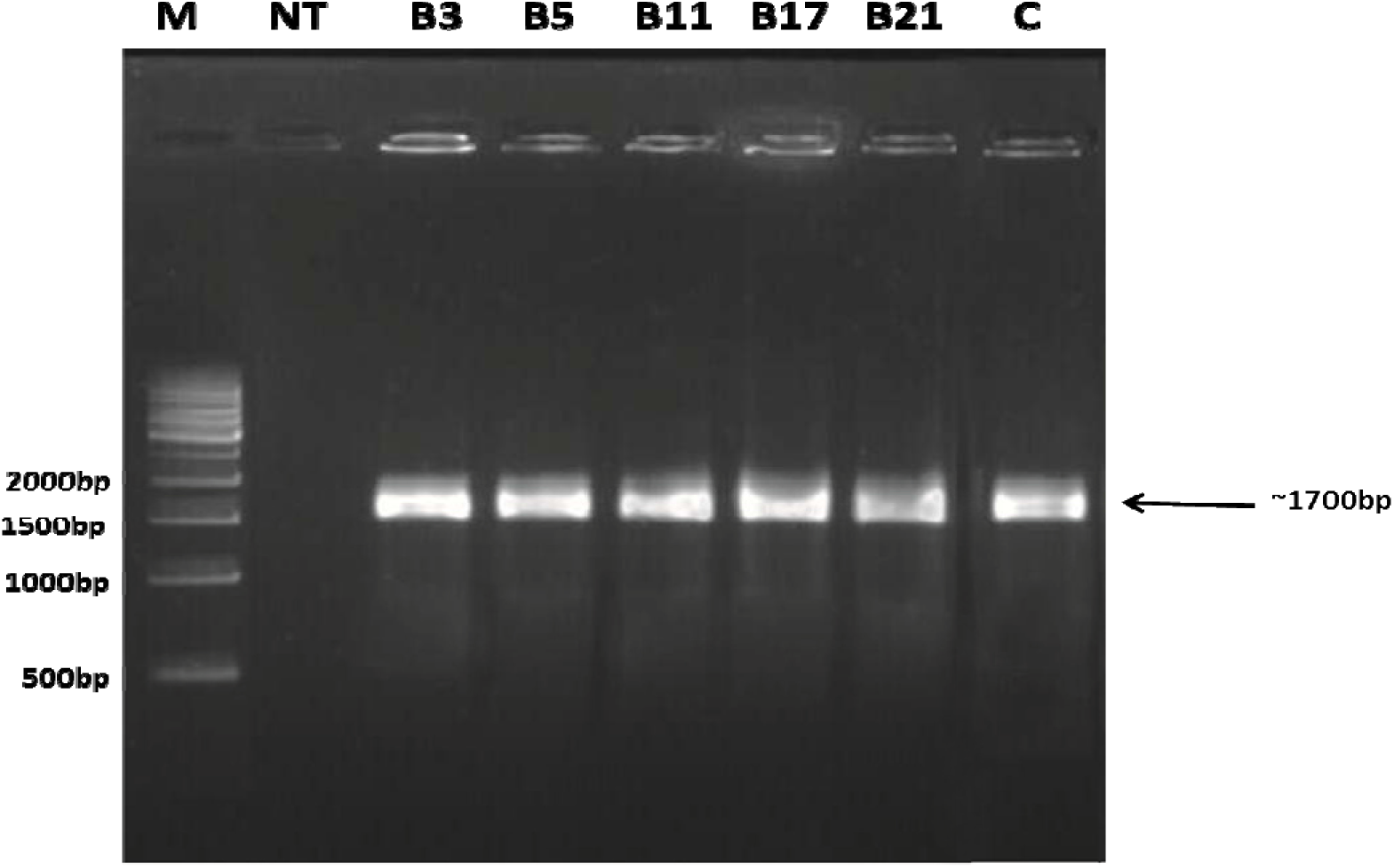
RT-PCR products amplified from the total RNA of transgenic plant. Primers designed for PCR amplification of the *cagA* gene which specifically amplified DNA fragments of ∼ 1700bp. Lane description: molecular-weight ladder (M), RT-PCR product of cDNA from a non-transformed plant (NT), RT-PCR products of cDNA from transformed plant lane No. B3, B5, B11, B17, B21. pBI121-*cagA* plasmid was used as positive control (C).

### Expression analysis of the cagA gene by real-time PCR

To determine the quality of the translation among the samples, the Amplicons from the real-time PCR were performed on a 2% agarose gel. HKG was found to be expressed at roughly 150bp in all of the samples. The *cagA* gene was found to be expressed in each of the five samples studied. Despite the fact that the expression was consistent and visible at around 151bp, the samples B3, B5, B11, and B17 were determined to be overexpressed compared to B21. The *cagA* gene was overexpressed in samples B11 and B17, with 17.3302 0.42466 and 21.2610 0.416804, respectively. B5 (11.59 0.284) was found to be underexpressed [Figure 4].

**Figure 4:**
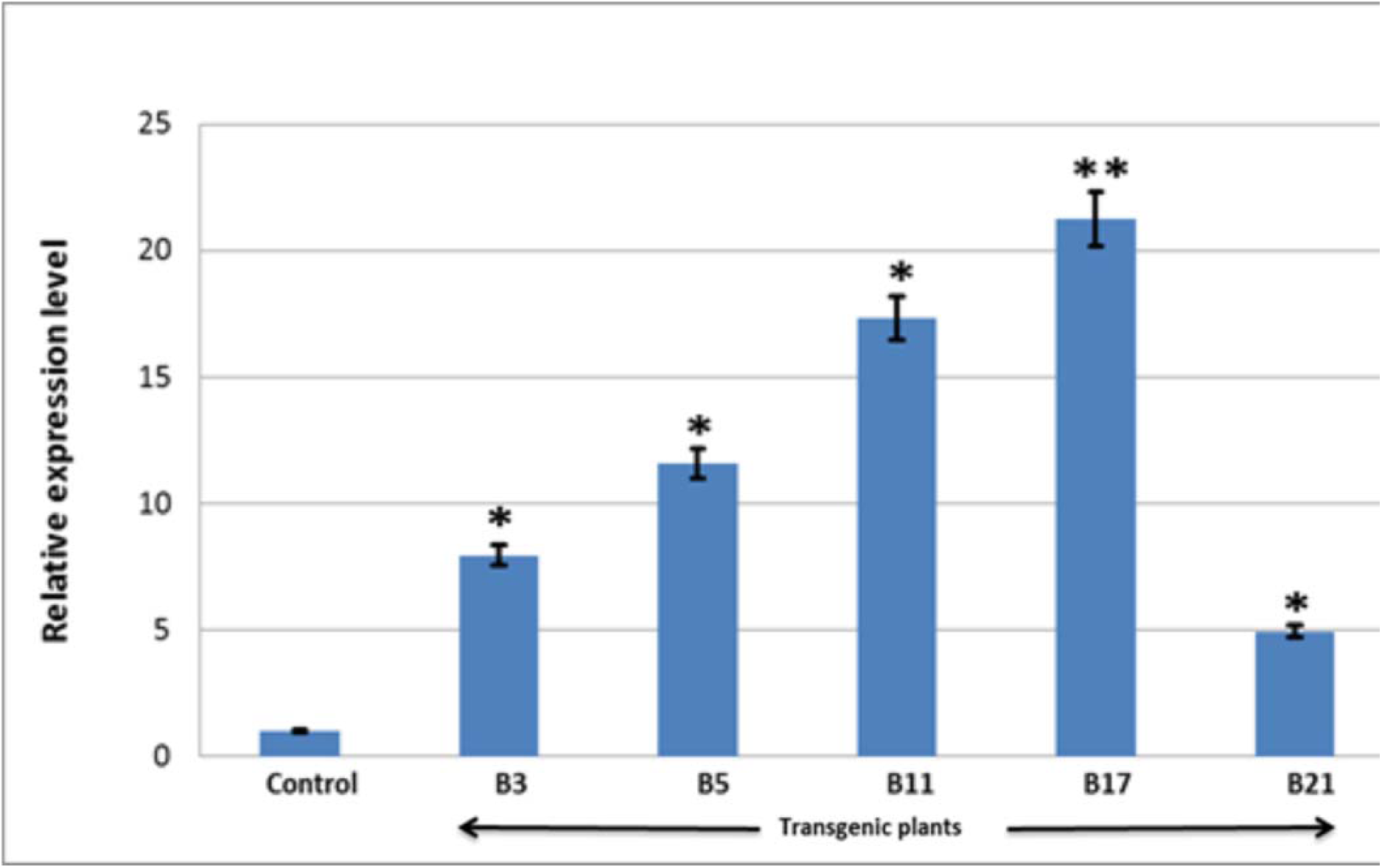
Relative expression levels of the *cagA* gene in transgenic plant brinjals. Graph showing the relative expression levels of *cagA* gene of the transgenic and control plants. Control expression was considered to be 1 and 100%. All the values are average of duplicates. Values are expressed as value ± SD. ** Highly significant; * Significant at 5% level.

These numbers correspond to the band expression on the gel. The F-value (0.003**) is significant, according to the results. As a result, no assumption of equal variance is made. We reject the null hypothesis since the p-value (0.009**) is significant at a 5% level, indicating that there is statistical significance in terms of relative expression levels between non-transgenic (control) and transgenic Brinjal. There is a statistically significant difference between the average relative expression level of Brinjal non-transgenic plants (control) and Brinjal transgenic plants. The following hypotheses were investigated using the t-test (paired t-test) in SPSS (Statistical Package for Social Science) on the differences between Brinjal non-transgenic plants (control) and Brinjal transgenic plants. At the 5% level, the t-value for all five pairings, B3 *cagA* – Control (0.025*) & B5 *cagA* – Control (0.012*) & B11 *cagA* – Control (0.012*) & B17 *cagA* – Control (0.009**) & B21 *cagA* – Control (0.041*), is significant. We reject the null hypothesis because the p-value is significant at 5%, indicating that there is a statistical difference verifying B3, B5, B11, B17, B21 relative expressions larger than or equal to Control [Figure 4].

### Expression analysis of cagA protein in transgenic brinjals

The study’s primary goal was to create *cagA* proteins in transgenic plants. Within transgenic brinjal plants, Western blotting indicated large levels of *cagA* protein at around 63.5 kDa [Figure 5]. There was no protein band at the appropriate molecular size on a blot of protein from non-transgenic or control mice. The transgenic brinjal lines B3, B5, B11, B17, and B21, on the other hand, showed a high positive connection between *cagA* transcripts and protein levels. In comparison to the other samples, the *cagA* protein was expressed more in the B11 and B17 samples. The RT-PCR results and the amount of *cagA* protein acquired from western blot analysis were determined to be in agreement. Bands of beta-actin were detected with a diameter of 42 kDa [Figure 5].

**Figure 5:**
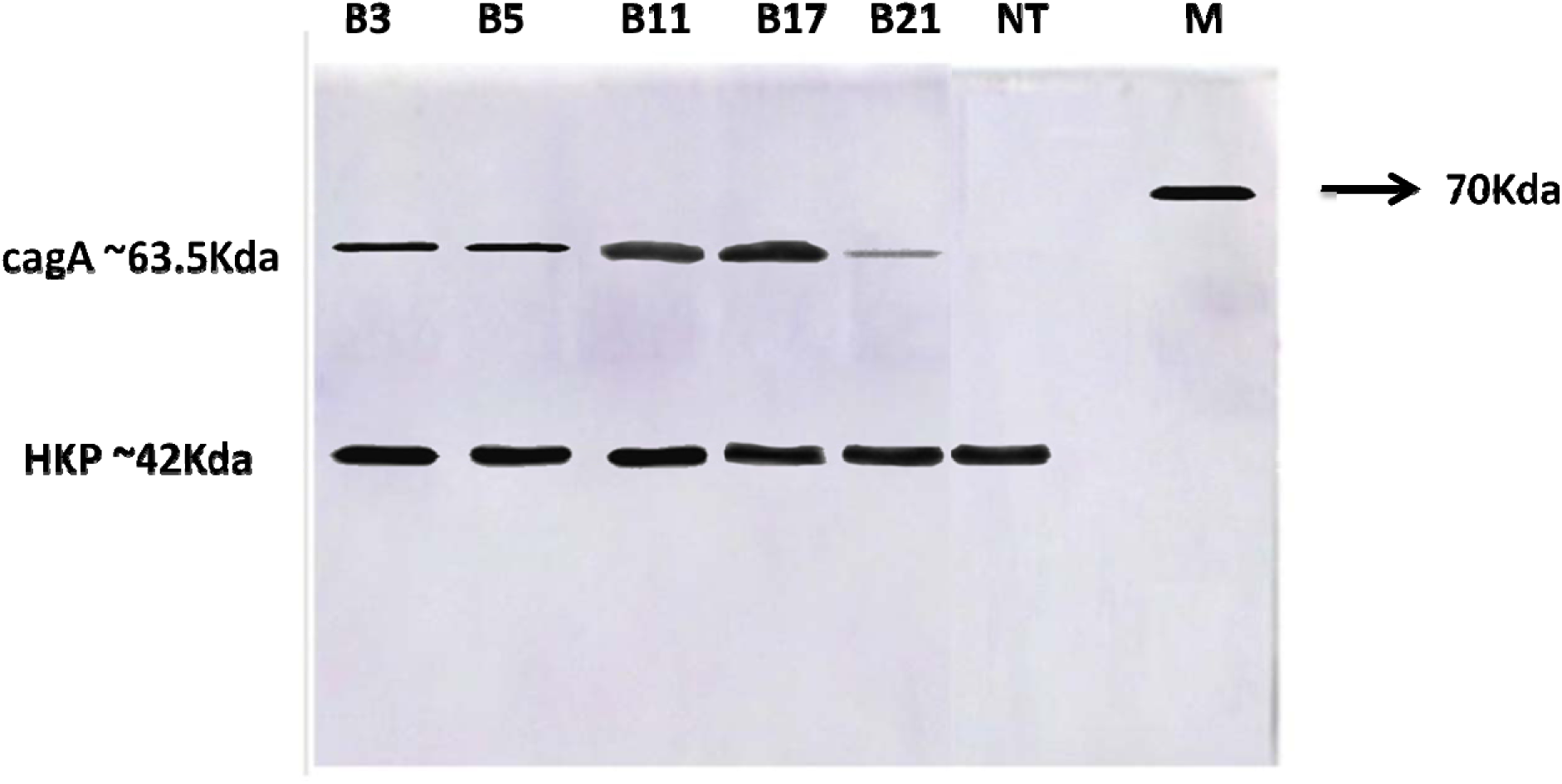
Western-blot analyses of transgenic brinjals. Western-blot analysis of *cagA* protein from independent transgenic plants. B3, B5, B11, B17, B21 independent transgenic brinjals and NT is the non-transgenic plant brinjal, Beta Actin was used as HKP. Bands seen at ∼63.5KDa corresponds to *cagA* protein and Beta Actin bands (HKP) were seen at ∼42KDa.

### Immunohistochemistry assay

Immunohistochemical analysis was used to examine the changes in *cagA* protein distribution inside the leaves of both transgenic and non-transgenic plants. The distribution of *cagA* in non-transgenics was scarcely apparent in the samples, whereas *cagA* immunohistochemistry specific staining was found in the parenchymal cells underneath the epidermal areas of the leaves in transgenics. The epidermal layer had very little specific signal, whereas the parenchymal sections had a lot. The signal appears to be entirely visible in the cytoplasm rather than in the cell walls. To our knowledge we feel this is the first time this study has been published. According to research published by Qing Gu et al., 2006, a systemic strain of *H. pylori*, ZJC02, produced a full-length *ureB* gene in transgenic rice plants, resulting in antigen-based protection (Gu et al., 2006). The immunohistochemistry findings obtained in our investigation were similar to those reported by Qing Gu et al, with the exception that our protein was *cagA* [Figure 6].

**Figure 6:**
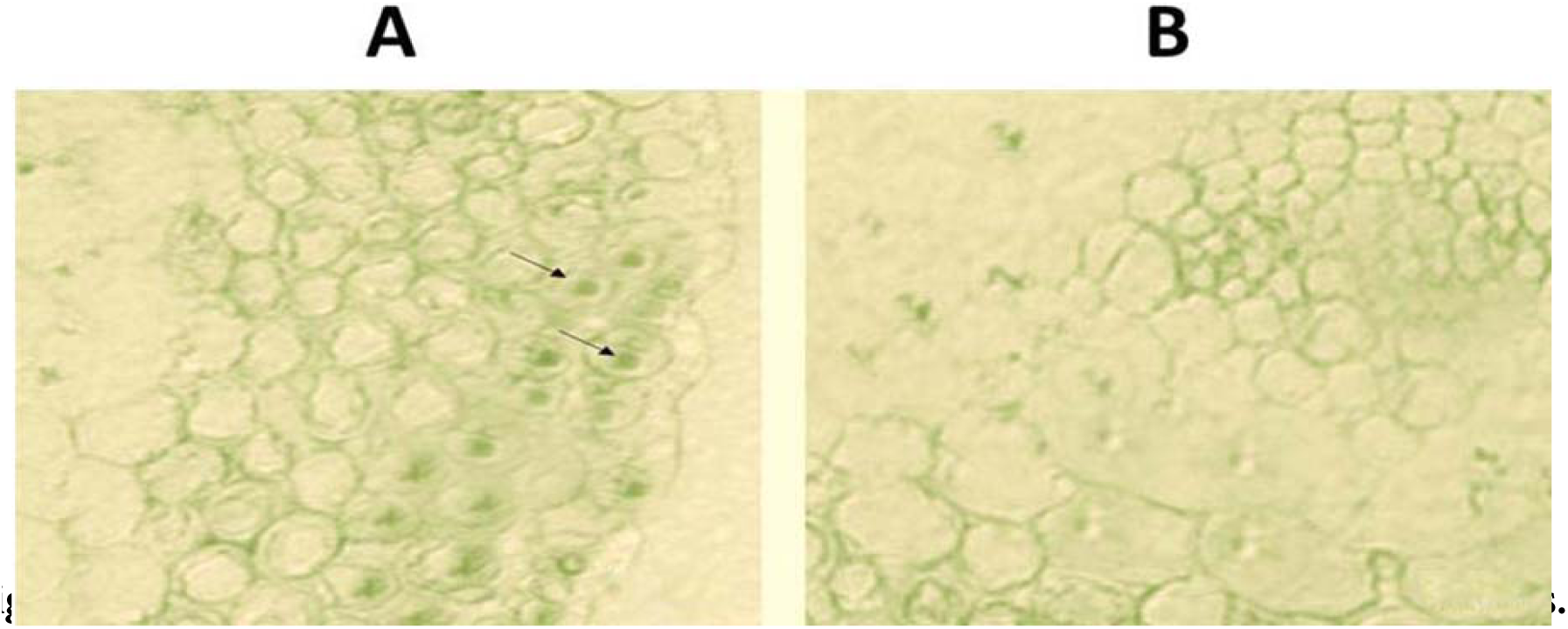
Images showing the distribution of *cagA* protein in Brinjal leaves. (**A):** Transgenics expressing the *cagA* protein as shown with arrows. (**B):** Non-transgenics without expression. Samples without antibodies and with PBS was used as negative control (not shown in the study). All the images shown here are represented from three independent cross-sections and three-leaf samples (Bars = 30μm).

### ELISA analysis

The ELISA test demonstrated that transgenic brinjals generate a large quantity of *cagA* protein. The spectrophotometric measurement of the diluted samples’ antibody concentrations was shown on a graph [Figure 7]. The largest quantity of *cagA* antigen between the transgenic brinjal lines was detected in B11 and B17 samples at both 1:5 and 1:10 dilutions, with the p-value significant at the 1% level, comparable to the RT-PCR and western blot. In the control sample, however, no antigen was found. The results of the RT-PCR and ELISA tests revealed a high positive connection between *cagA* transcripts and protein levels in transgenic brinjal.

**Figure 7:**
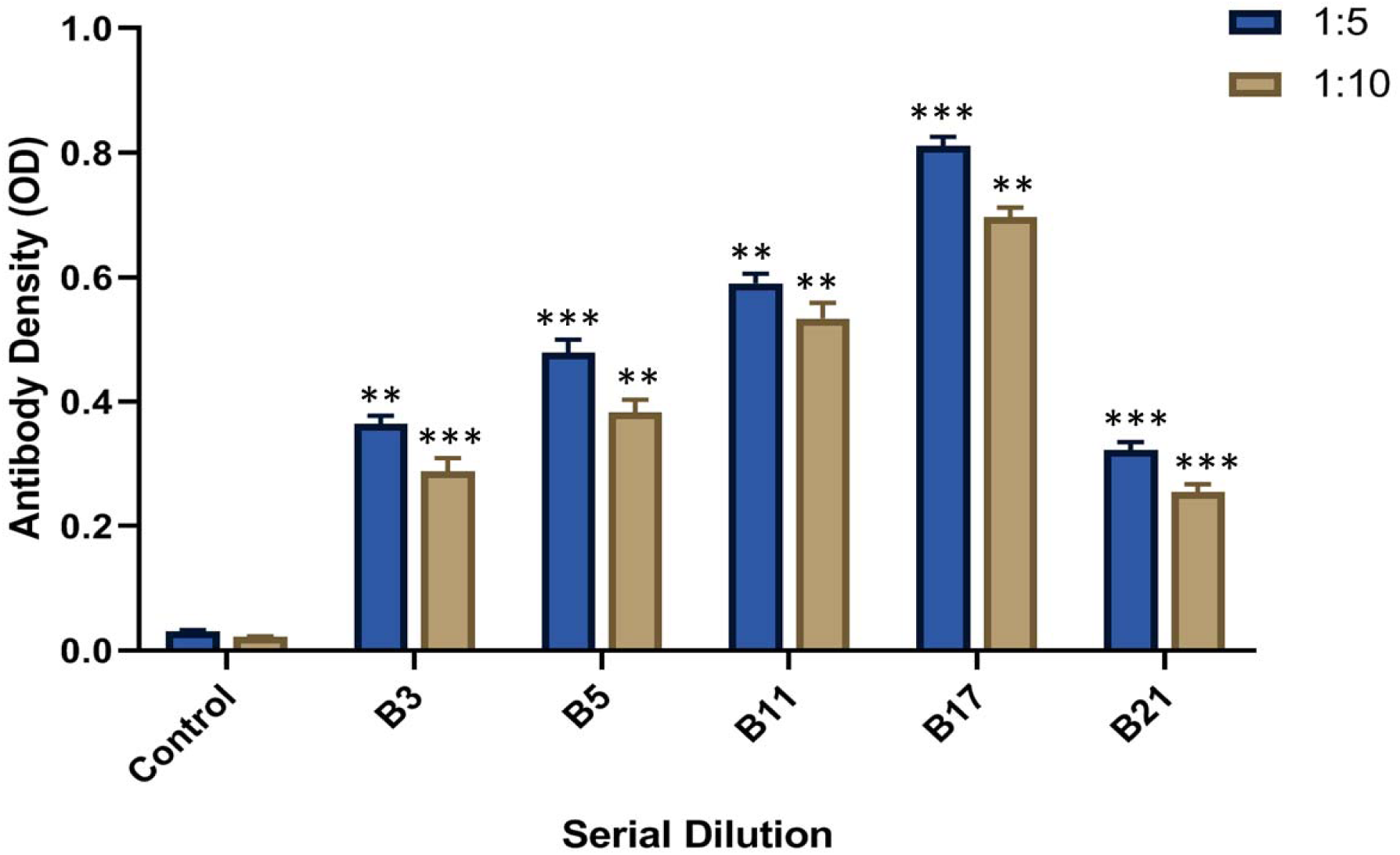
Data graph obtained from ELISA reader shows that the different amount of *cagA* antigen in transgenic brinjals and control plant with respect to 1/5 and 1/10 dilution ratios. No signal was observed in the control plant. All the values are average of triplicates. Values are expressed as value ± SD. *** Highly significant; ** Significant at 1% level.

## Discussion

The production of recombinant proteins within plants has been achieved with a vast number of expression systems (Mehran et al., 2014a; Mehran et al., 2014b). However, there are a number of issues that still rate limit the routine use of this technique to us plants as factories to produce recombinant biopharmaceuticals including vaccines. There is need to optimize a number of factors to make it most cost-effective. *Helicobacter pylori,* a worldwide prevalent bacterial strain, is stated to span about 40–50% of people globally (Nešić et al., 2014) and is usually associated with ulcers and cancers of the stomach, intestine and mucosa-associated lymphoid tissue (MALT) . This pathogen is known to colonize the stomach and usually accompanies chronic gastritis and other ulcers, which are proved to be risky to develop adenocarcinoma and lymphoma (Lyon, 1994).

Vaccines improve immunity against specific diseases and play vital role in managing disease pandemics. Vaccine is a protein that mimics the pathogen made by heat killing or weakening of the strain (Kurup and Thomas, 2020). They aid the immune system to detect the foreign antigen and subsequently act on it. While there are a number of approaches to produce vaccines, the plant-based vaccines known to reduce the concerns of conventional vaccines. Furthermore, the cost of plant based vaccine production less and the number of interventions are minimum (Zendehbad et al., 2014; Ma et al., 2020). Expressing biopharmaceutical proteins within the transgenic plants is a cost-effective mode to reduce the production costs and other limitations associated with the expression systems (Yao et al., 2015). Plant based vaccines are modified genetically and gene-encoding pathogens (bacteria or virus) can be directly inserted into plants without fading off their immunogenicity (Stern and Markel, 2005). In this present study, *cagA* protein was transgenically produced in brinjal plants. This is the first report regarding the expression of *cagA* gene in brinjal plants. Our experimental data demonstrated that *cagA* protein could be expressed in transgenic brinjal plants. The expression of the *cagA* gene in brinjal plants was verified by RT-PCR and protein electrophoresis. Our PCR amplification and blotting methods also confirmed the expression of *cagA* gene. Further, the western blot analysis revealed the accumulation of *cagA* protein in the tissue, which could be the vaccine of choice. Previously, *Helicobacter pylori ureB* antigen gene was also expressed into the rice genome using *Agrobacterium*-mediated transformation and a set of 30 regenerated plants were reported to occumulate *ureB* antigen, which was confirmed by PCR and blotting (Gu et al., 2006).

Antibiotics resistance in strains is a major problem, hence, developing a new vaccine is the need of the hour (Yang et al., 2011). In both developing and developed countries, *H. pylori*, has high infection rates and also gained multiple antibiotic resistance (Schillberg et al., 2003). Even then, the cost-effectiveness of the vaccination has remained an important area of research. Even though many of the eukaryotic proteins can be produced at a low cost and in a more viable manner, most of the post-translational modifications cannot occur in prokaryotes, making them inefficient (Giddings et al., 2000).

The *cagA* gene was successfully cloned into the binary vector pBI121 using kanamycin resistance gene as marker. The gene was successfully cloned and transformed into the brinjal callus through *Agrobacterium*-mediated method. Out of the 52 plants, a set five plants were found to be transgenic based on presence of the *cagA* gene product. Expression of the *cagA* gene at mRNA level in five plants was confirmed by RT-PCR and translation by Western blot analysis. The outturn of these experiments suggests that the *cagA* transgenic brinjal can further be evaluated to be used as a vaccine candidate against *H.pylori*.

Edible vaccines are produced by inserting the desired transgene into the selected plant cell. In the current scenario, edible vaccines are developed for both veterinary and human use. The main challenge lies in convincing people to use such vaccines. Moreover, these edible vaccines are economical and safe when compared to other conventional vaccines available in the market (Kalbina et al., 2010). In conclusion, our results demonstrated the successful integration of the desired *cagA* gene into the nuclear genome of the transgenic brinjal and can serve as potential candidate events for prospecting the purification and use as vaccine.

## Declarations

## Acknowledgements

The authors acknowledge the supports from JSS Research Foundation, JSS Technical Institutions Campus, Mysore; Genei Laboratories Pvt Ltd, Bengaluru, Department of Biotechnology and Crop Improvement, Postgraduate Centre, College of Horticulture, University of Horticulture Sciences Campus, GKVK, Bengaluru and Postgraduate Department of Biotechnology, JSS College, Mysore.

## Availability of data and material

All the data and supplementary materials are available from the corresponding author on reasonable request.

## Consent for publication

All authors consent to publish this work.

## Ethics approval

The experimentation was conducted with prior approval from the Institutional Biosafety Committee (IBSC) of JSS College, Mysore at Postgraduate Department of Biotechnology, JSS College, Mysuru, Genei India Pvt. Ltd., Bangalore and Department of Biotechnology and Crop Improvement, College of Horticulture, University of Horticulture Sciences, GKVK Post, Bangalore (IBSC Registration number: JSSC111220191080). Appropriate permission for the collection of plant or seed specimens has been obtained from College of Horticulture, University of Horticulture Sciences, GKVK Post, Bangalore.

## References

1. Abdoli-Nasab, M., Jalali-Javaran, M., Cusidó, R.M., Palazón, J., Baghizadeh, A. and Alizadeh, H., 2013. Expression of the truncated tissue plasminogen activator (K2S) gene in tobacco chloroplast. Molecular biology reports 40, 5749–5758.

2. Alaguponniah, S., Krishna, D.V., Paul, S., Christyraj, J.R.S.S., Nallaperumal, K. and Sivasubramaniam, S., 2020. Finding of novel telomeric repeats and their distribution in the human genome. Genomics 112, 3565–3570. Available from: https://doi.org/10.1016/j.ygeno.2020.04.010

3. Arumugaperumal, A., Paul, S., Lathakumari, S., Balasubramani, R. and Sivasubramaniam, S., 2020. The draft genome of a new Verminephrobacter eiseniae strain: a nephridial symbiont of earthworms. Annals of microbiology 70, 1–18. Available from: https://doi.org/10.1186/s13213-020-01549-w

4. Backert, S., Clyne, M. and Tegtmeyer, N., 2011. Molecular mechanisms of gastric epithelial cell adhesion and injection of CagA by Helicobacter pylori. Cell Communication and Signaling 9, 1–11.

5. BARZIGAR, R., HARAPRASADN, K.B. and Mehran, M., 2020. Challenges and recent developments associated with vaccine antigens production against Helicobacter pylori. Int J App Pharm 12, 45–50.

6. Barzigar, R., Haraprasad, N., Kumar, B.Y.S. and Mehran, M.J., 2021. Cloning and Expression of Vacuolating Cytotoxin A (VacA) Antigenic Protein in Nicotiana benthamiana Leaves a Potential Source of the Vaccine against Helicobacter pylori. International Journal of Pharmaceutical Investigation 11.

7. Barzigar, R., Mehran, M.J., Haraprasad, N., Kumar, B.Y.S. and Fakrudin, B., 2022. Transient recombinant expression of highly immunogenic CagA, VacA and NapA in Nicotiana benthamiana. Biotechnology Reports 33, e00699.

8. Bleve, G., Rizzotti, L., Dellaglio, F. and Torriani, S., 2003. Development of reverse transcription (RT)-PCR and real-time RT-PCR assays for rapid detection and quantification of viable yeasts and molds contaminating yogurts and pasteurized food products. Applied and environmental microbiology 69, 4116–4122.

9. Chatterjee, A., Paul, S., Bisht, B., Bhattacharya, S., Sivasubramaniam, S. and Paul, M.K., 2021. Advances in targeting the WNT/β-catenin signaling pathway in cancer. Drug Discovery Today. Available from: https://doi.org/10.1016/j.drudis.2021.07.007

10. Chinnadurai, L., Eswaramoorthy, T., Paramachandran, A., Paul, S., Rathy, R., Arumugaperumal, A., Sivasubramaniam, S. and Thavasimuthu, C., 2018. Draft genome sequence of Escherichia coli phage CMSTMSU, isolated from shrimp farm effluent water. Microbiology Resource Announcements 7. Available from: https://doi.org/10.1128/MRA.01034-18

11. Expósito-Rodríguez, M., Borges, A.A., Borges-Pérez, A. and Pérez, J.A., 2008. Selection of internal control genes for quantitative real-time RT-PCR studies during tomato development process. BMC plant biology 8, 1–12.

12. Fallone, C.A., Chiba, N., van Zanten, S.V., Fischbach, L., Gisbert, J.P., Hunt, R.H., Jones, N.L., Render, C., Leontiadis, G.I. and Moayyedi, P., 2016. The Toronto consensus for the treatment of Helicobacter pylori infection in adults. Gastroenterology 151, 51–69. e14.

13. Garcia Jr G., Paul, S., Beshara, S., Ramanujan, V.K., Ramaiah, A., Nielsen-Saines, K., Li, M.M., French, S.W., Morizono, K. and Kumar, A., 2020. Hippo signaling pathway has a critical role in Zika virus replication and in the pathogenesis of neuroinflammation. The American Journal of Pathology 190, 844–861. Available from: https://doi.org/10.1016/j.ajpath.2019.12.005

14. Giddings, G., Allison, G., Brooks, D. and Carter, A., 2000. Transgenic plants as factories for biopharmaceuticals. Nature biotechnology 18, 1151–1155.

15. Graham, D.Y. and Shiotani, A., 2008. New concepts of resistance in the treatment of Helicobacter pylori infections. Nature Clinical Practice Gastroenterology & Hepatology 5, 321–331.

16. Green, M.R. and Sambrook, J., 2019. Amplification of cDNA generated by reverse transcription of mRNA: two-step reverse transcription-polymerase chain reaction (RT-PCR). Cold Spring Harbor Protocols 2019, pdb. prot095190.

17. Gu, Q., Han, N., Liu, J. and Zhu, M., 2006. Expression of Helicobacter pylori urease subunit B gene in transgenic rice. Biotechnology letters 28, 1661–1666.

18. Hatakeyama, M., 2017. Structure and function of Helicobacter pylori CagA, the first-identified bacterial protein involved in human cancer. Proceedings of the Japan Academy, Series B 93, 196–219.

19. Hooi, J.K., Lai, W.Y., Ng, W.K., Suen, M.M., Underwood, F.E., Tanyingoh, D., Malfertheiner, P., Graham, D.Y., Wong, V.W. and Wu, J.C., 2017. Global prevalence of Helicobacter pylori infection: systematic review and meta-analysis. Gastroenterology 153, 420–429.

20. Iannacone, R., Grieco, P.D. and Cellini, F., 1997. Specific sequence modifications of a cry3B endotoxin gene result in high levels of expression and insect resistance. Plant molecular biology 34, 485–496.

21. Josephson, M. and Skole, K., 2018. The Houston Consensus Conference on Testing for Helicobacter pylori infection. Clinical Gastroenterology and Hepatology 16, 2004–2005.

22. Kalbina, I., Engstrand, L., Andersson, S. and Strid, Å., 2010. Expression of Helicobacter pylori TonB protein in transgenic Arabidopsis thaliana: toward production of vaccine antigens in plants. Helicobacter 15, 430–437.

23. Kumar, A., Garcia Jr, G., Paul, S., Morizono, K. and Arumugaswami, V., 2020. Hippo Signaling Pathway has a critical role in Zika Virus Replication and in the Pathogenesis of neuroinflammation. Investigative Ophthalmology & Visual Science 61, 440–440. Available from: https://iovs.arvojournals.org/article.aspx?articleid=2766536

24. Kumavath, R., Paul, S., Pavithran, H., Paul, M.K., Ghosh, P., Barh, D. and Azevedo, V., 2021. Emergence of Cardiac Glycosides as Potential Drugs: Current and Future Scope for Cancer Therapeutics. Biomolecules 11, 1275. Available from: https://doi.org/10.3390/biom11091275

25. Kurup, V.M. and Thomas, J., 2020. Edible vaccines: promises and challenges. Molecular biotechnology 62, 79–90.

26. Li, Z., Parris, S. and Saski, C.A., 2020. A simple plant high-molecular-weight DNA extraction method suitable for single-molecule technologies. Plant methods 16, 1–6.

27. Livak, K.J. and Schmittgen, T.D., 2001. Analysis of relative gene expression data using real-time quantitative PCR and the 2− ΔΔCT method. methods 25, 402–408.

28. Lyon, F., 1994. IARC monographs on the evaluation of carcinogenic risks to humans. Some industrial chemicals 60, 389–433.

29. Ma, Y., Lee, C.-J. and Park, J.-S., 2020. Strategies for Optimizing the Production of Proteins and Peptides with Multiple Disulfide Bonds. Antibiotics 9, 541.

30. Malfertheiner, P., Megraud, F., O’morain, C., Gisbert, J., Kuipers, E., Axon, A., Bazzoli, F., Gasbarrini, A., Atherton, J. and Graham, D.Y., 2017. Management of Helicobacter pylori infection—the Maastricht V/Florence consensus report. Gut 66, 6–30.

31. Mehran, M.J., Kumar, B.Y.S., Haraprasad, N., Barzigar, R., Fakrudin, B. and Paul, S. Evaluation of iceA1 Gene Expression of Helicobacter pylori Risk Factor of Gastric Cancer in Transgenic Brinjal. Available from: https://ijper.org/article/1580

32. Mehran, M.J., Kumar, B.Y.S., Haraprasad, N., Barzigar, R. and Paul, S., 2021. Cloning and Expression of Helicobacter pylori ulcer Associated Gene-iceA1 in Brinjal (Solanum melongena L.). International Journal of Pharmaceutical Investigation 11, 338–344. Available from: https://www.jpionline.org/index.php/ijpi/article/view/1189/610

33. Mehran, M.J., Zendehbad, S.H. and Malla, S., 2014a. Cloning and expression of a partial UreA antigen for the production of vaccine against helicobacter pylori, the risk factor for gastric cancer. Asian J Pharm Clin Res 7, 111–117.

34. Mehran, M.J., Zendehbad, S.H. and Malla, S., 2014b. Free radical scavenging and antioxidant potential activity of cassava plants. Asian J Pharm Clin Res 7, 66–70.

35. Meyer, R.S., Karol, K.G., Little, D.P., Nee, M.H. and Litt, A., 2012. Phylogeographic relationships among Asian eggplants and new perspectives on eggplant domestication. Molecular phylogenetics and evolution 63, 685–701.

36. Miao, X., Kumar, R.R., Shen, Q., Wang, Z., Zhao, Q., Singh, J., Paul, S., Wang, W. and Shang, X., 2022. Phytoremediation for Co-contaminated Soils of Cadmium and Polychlorinated Biphenyls Using the Ornamental Plant Tagetes patula L. Bulletin of Environmental Contamination and Toxicology, 1–7. Available from: https://doi.org/10.1007/s00128-021-03392-4

38. Nayeri, F.D. and Anbuhi, M.H., 2019. Transient expression of etanercept therapeutic protein in tobacco (Nicotiana tabacum L.). International journal of biological macromolecules 130, 483–490.

39. Nešić, D., Buti, L., Lu, X. and Stebbins, C.E., 2014. Structure of the Helicobacter pylori CagA oncoprotein bound to the human tumor suppressor ASPP2. Proceedings of the National Academy of Sciences 111, 1562–1567.

40. Ohnishi, N., Yuasa, H., Tanaka, S., Sawa, H., Miura, M., Matsui, A., Higashi, H., Musashi, M., Iwabuchi, K. and Suzuki, M., 2008. Transgenic expression of Helicobacter pylori CagA induces gastrointestinal and hematopoietic neoplasms in mouse. Proceedings of the National Academy of Sciences 105, 1003–1008.

41. Pan, X., Ke, H., Niu, X., Li, S., Lv, J. and Pan, L., 2018. Protection against Helicobacter pylori infection in BALB/c mouse model by oral administration of multivalent epitope-based vaccine of cholera toxin B subunit-HUUC. Frontiers in immunology 9, 1003.

42. Patel, M., Dewey, R.E. and Qu, R., 2013. Enhancing Agrobacterium tumefaciens-mediated transformation efficiency of perennial ryegrass and rice using heat and high maltose treatments during bacterial infection. Plant cell, tissue and organ culture (PCTOC) 114, 19–29.

43. Paul, S., Arumugaperumal, A., Balakrishnan, S., Sathiya Balasingh Thangapandi, E.J.J., Sarjubala Devi, H., Johnson, T., Maisnam, S., Soman Syamala, S., Ramamoorthy, S. and Karthikeyan, R., 2023. PATHOGENICITY, STRUCTURE AND GENOME ANALYSIS OF THE GRANULOVIRUS INFECTING THE CATERPILLARS OF PIERIS BRASSICAE LINN. (INSECTA: LEPIDOPTERA: PIERIDAE). J. Exp. Zool. India 26, 1219–1240. Available from: https://doi.org/10.51470/jez.2023.26.1.1219

44. Paul, S., Balakrishnan, S., Arumugaperumal, A., Lathakumari, S., Syamala, S.S., Arumugaswami, V. and Sivasubramaniam, S., 2021a. The transcriptome of anterior regeneration in earthworm Eudrilus eugeniae. Molecular Biology Reports 48, 259–283. Available from: https://doi.org/10.1007/s11033-020-06044-8

45. Paul, S., Balakrishnan, S., Arumugaperumal, A., Lathakumari, S., Syamala, S.S., Vijayan, V., Durairaj, S.C.J., Arumugaswami, V. and Sivasubramaniam, S., 2022a. Importance of clitellar tissue in the regeneration ability of earthworm Eudrilus eugeniae. Functional & Integrative Genomics, 1–32. Available from: https://doi.org/10.1007/s10142-022-00849-5

46. Paul, S., Balakrishnan, S., Arumugaperumal, A. and Thangapandi EJJSB, D.H., 2021b. Genome sequencing, Analysis and Characterization of Baculovirus Infecting the Caterpillar, Spilosoma Obliqua Walker (Arctiidae)(Insecta: Lepidoptera) from India. J Virol Antivir Res 10 5, 2. Available from: https://www.scitechnol.com/peer-review/genome-sequencing-analysis-and-characterization-of-baculovirus-infecting-the-caterpillar-spilosoma-obliqua-walker-arctiidae-insect-Gi2Y.php?article_id=17284

47. Paul, S., Heckmann, L.-H., Sørensen, J.G., Holmstrup, M., Arumugaperumal, A. and Sivasubramaniam, S., 2018. Transcriptome sequencing, de novo assembly and annotation of the freeze tolerant earthworm, Dendrobaena octaedra. Gene Reports 13, 180–191. Available from: https://doi.org/10.1016/j.genrep.2018.10.010

48. Paul, S., Sudalai Mani, D.K., Sandhya, S.S., Subburathinam, B., Vijithkumar, V., Vaithilingaraja, A. and Sudhakar, S., 2022b. Identification, tissue specific expression analysis and functional characterization of arrestin gene (ARRDC) in the earthworm Eudrilus eugeniae: a molecular hypothesis behind worm photoreception. Molecular Biology Reports, 1–12. Available from: https://doi.org/10.1007/s11033-022-07256-w

49. Ponesakki, V., Syamala, S.S., Paul, S., Arumugaperumal, A., Selvakumar, P., Vijayan, V., Raj, A.P.M.S. and Sivasubramaniam, S., 2023. ACTIVATION OF MOUSE EAR LOBE TISSUE REGENERATION BY METABOLITES OF EARTHWORM. J. Exp. Zool 26, 1191–1205. Available from: https://doi.org/10.51470/jez.2023.26.1.1191

50. Ramalho, A.S., Beck, S., Farinha, C.M., Clarke, L.A., Heda, G.D., Steiner, B., Sanz, J., Gallati, S., Amaral, M.D. and Harris, A., 2004. Methods for RNA extraction, cDNA preparation and analysis of CFTR transcripts. Journal of Cystic Fibrosis 3, 11–15.

51. Ramasamy Rajesh Kumar, S.A.D., Rashmi Saini, Punita Kumari, Priyanka Roy, Sayan Paul, 2020. Impacts on dietary habits and health of Indian population during COVID-19 lockdown. Public Health Review: International Journal of Public Health Research 7, 38–50. Available from: https://doi.org/10.17511/ijphr.2020.i06.01

52. Rana, I., Kataria, S., Tan, T.L., Hajam, E.Y., Kashyap, D.K., Saha, D., Ajnabi, J., Paul, S., Jayappa, S. and Ananthan, A.S., 2022a. MINDIN (SPONDIN-2) IS ESSENTIAL FOR CUTANEOUS FIBROGENESIS IN A MOUSE MODEL OF SYSTEMIC SCLEROSIS. Journal of Investigative Dermatology 143, 699–710. Available from: https://doi.org/10.1016/j.jid.2022.10.011

53. Rana, I., Kataria, S., Tan, T.L., Hajam, E.Y., Kashyap, D.K., Saha, D., Ajnabi, J., Paul, S., Jayappa, S. and Ananthan, A.S., 2022b. Mindin is essential for cutaneous fibrogenesis in a new mouse model of systemic sclerosis. bioRxiv, 2022.01. 26.477822. Available from: https://doi.org/10.1101/2022.01.26.477822

54. Sabalza, M., Christou, P. and Capell, T., 2014. Recombinant plant-derived pharmaceutical proteins: current technical and economic bottlenecks. Biotechnology letters 36, 2367–2379.

55. Saini, D.K. and Kaushik, P., 2019. Visiting eggplant from a biotechnological perspective: A review. Scientia Horticulturae 253, 327–340.

56. Sambrook, J. and Russell, D.W., 2006. Preparation of plasmid DNA by alkaline lysis with SDS: minipreparation. Cold Spring Harbor Protocols 2006, pdb. prot4084.

57. Saravanakumari, A., Selvi, D., Rajesh Kumar, R. and Paul, S., 2020. Assessment of patient satisfaction with inpatient services at secondary level setting. Available from: https://doi.org/10.17511/ijphr.2020.i06.03

58. Schillberg, S., Fischer, R. and Emans, N., 2003. Molecular farming of recombinant antibodies in plants. Cellular and Molecular Life Sciences CMLS 60, 433–445.

59. Schmidt, G.W. and Delaney, S.K., 2010. Stable internal reference genes for normalization of real-time RT-PCR in tobacco (Nicotiana tabacum) during development and abiotic stress. Molecular Genetics and Genomics 283, 233–241.

60. Soria-Guerra, R.E., Rosales-Mendoza, S., Márquez-Mercado, C., López-Revilla, R., Castillo-Collazo, R. and Alpuche-Solís, Á.G., 2007. Transgenic tomatoes express an antigenic polypeptide containing epitopes of the diphtheria, pertussis and tetanus exotoxins, encoded by a synthetic gene. Plant cell reports 26, 961–968.

61. Stern, A.M. and Markel, H., 2005. The history of vaccines and immunization: familiar patterns, new challenges. Health affairs 24, 611–621.

62. Streatfield, S.J. and Howard, J.A., 2003. Plant production systems for vaccines. Expert review of Vaccines 2, 763–775.

63. Tatta, E.R., Paul, S. and Kumavath, R., 2023. Transcriptome Analysis revealed the Synergism of Novel Rhodethrin inhibition on Biofilm architecture, Antibiotic Resistance and Quorum sensing in Enterococcus faecalis. Gene 871, 147436. Available from: https://doi.org/10.1016/j.gene.2023.147436

64. Tiwari, S., Verma, P.C., Singh, P.K. and Tuli, R., 2009. Plants as bioreactors for the production of vaccine antigens. Biotechnology advances 27, 449–467.

65. Turabi, K.S., Deshmukh, A., Paul, S., Swami, D., Siddiqui, S., Kumar, U., Naikar, S., Devarajan, S., Basu, S. and Paul, M.K., 2022. Drug repurposing—an emerging strategy in cancer therapeutics. Naunyn-Schmiedeberg’s Archives of Pharmacology 395, 1139–1158. Available from: https://doi.org/10.1007/s00210-022-02263-x

66. Untergasser, A., Cutcutache, I., Koressaar, T., Ye, J., Faircloth, B.C., Remm, M. and Rozen, S.G., 2012. Primer3—new capabilities and interfaces. Nucleic acids research 40, e115–e115.

67. Wang, W., Scali, M., Vignani, R., Spadafora, A., Sensi, E., Mazzuca, S. and Cresti, M., 2003. Protein extraction for two-dimensional electrophoresis from olive leaf, a plant tissue containing high levels of interfering compounds. Electrophoresis 24, 2369–2375.

68. Wei, Q., Du, L., Wang, W., Hu, T., Hu, H., Wang, J., David, K. and Bao, C., 2019. Comparative transcriptome analysis in eggplant reveals selection trends during eggplant domestication. International journal of genomics 2019.

69. Yang, C.-y., Chen, S.-y. and Duan, G.-c., 2011. Transgenic peanut (Arachis hypogaea L.) expressing the urease subunit B gene of Helicobacter pylori. Current microbiology 63, 387–391.

70. Yao, J., Weng, Y., Dickey, A. and Wang, K.Y., 2015. Plants as factories for human pharmaceuticals: applications and challenges. International journal of molecular sciences 16, 28549–28565.

71. Zendehbad, S.H., Mehran, M.J. and Malla, S., 2014. Flavonoids and phenolic content in wheat grass plant (Triticum aestivum). Asian J Pharm Clin Res 7, 184–187.

